# Villification of the intestinal epithelium is driven by Foxl1

**DOI:** 10.1101/2024.02.27.582300

**Authors:** Guoli Zhu, Deeksha Lahori, Jonathan Schug, Klaus H Kaestner

## Abstract

The primitive gut tube of mammals initially forms as a simple cylinder consisting of the endoderm-derived, pseudostratified epithelium and the mesoderm-derived surrounding mesenchyme. During mid-gestation a dramatic transformation occurs in which the epithelium is both restructured into its final cuboidal form and simultaneously folded and refolded to create intestinal villi and intervillus regions, the incipient crypts. Here we show that the mesenchymal winged helix transcription factor Foxl1, itself induced by epithelial hedgehog signaling, controls villification by activating BMP and PDGFRα as well as planar cell polarity genes in epithelial-adjacent telocyte progenitors, both directly and in a feed-forward loop with Foxo3.

## Introduction

The surface area of the human small intestine measures about 30 m^2^, or the size of half a badminton court (Helander and Fandriks 2014). This large area is required to enable effective digestion and absorption of nutrients. The major factor responsible for increasing the size of the intestinal epithelium is villification, or the formation of intestinal villi, which increase the surface area compared to a flat epithelium by close to 100-fold (Helander and Fandriks 2014). Remarkably, while villi are a feature of the small intestine in mammals and birds, they arise by divergent morphogenetic mechanisms. Thus, villus formation in birds is dependent on the sequential formation of first inner circular, then longitudinal, and finally muscularis mucosa muscles which coincide with the appearance of epithelial ridges, zigzags, and villi, respectively (Walton et al. 2018). In contrast, in mammals, villus formation can proceed in the absence of tensile forces generated by intestinal muscles; rather, it is dependent on PDGFRα/BMP positive mesenchymal cell clusters (Karlsson et al. 2000), which themselves are induced by epithelial hedgehog signaling (reviewed in (Walton et al. 2016a).

At the onset of villification, also termed ‘epithelial transition’, the endoderm-derived epithelium forms a simple tube of pseudostratified cells with a layer thickness of ∼ 50 µm. This pseudostratified epithelium, characterized by cell cycle-dependent interkinetic nuclear migration (Grosse et al. 2011), is then changed to a columnar epithelium coincident with villus formation. BMP signals from villus cluster mesenchymal cells restrict proliferation in the overlying epithelium so that cycling cells become restricted to intervillus regions, the precursors of the small intestinal crypts present in the adult. The critical role of BMP signaling in villus cluster formation was established by explant cultures of presumptive small intestine from 13.5 dpc (days post conception) embryos treated with localized sources of BMP or the pan-BMP inhibitor dorsomorphin (Walton et al. 2016b).

The induction of villus formation and the formation of villus clusters in the underlying mesenchyme are dependent on epithelial to mesenchymal hedgehog signaling. Thus, both Sonic Hedgehog (shh) and Indian Hedgehog (Ihh) are expressed in the epithelium, while the hedgehog receptors (Ptch1 and Ptch2) as well as the downstream transcription factor Gli1 are expressed in the mesenchyme (Motoyama et al. 1998; Ramalho-Santos et al. 2000). Because the two epithelial hedgehog proteins are partially redundant, Madison and colleagues addressed the role of hedgehog signaling in the developing intestine using epithelial expression of the pan-hedgehog inhibitor Hhip (Madison et al. 2005). Suppression of hedgehog signaling impaired villus formation, which was partially due to decreased BMP expression in the underlying mesenchyme.

More than 25 years ago, the winged helix transcription factor Foxl1 (formerly termed *Fhh6*) was shown to be expressed in the first one to two cell layers of the mesenchyme juxtaposed to the epithelium before villus formation by mRNA in situ hybridization (Kaestner et al. 1997), which was recently confirmed by immunofluorescence staining (Kondo and Kaestner 2021). This expression persists in adult mice and defines a subpopulation of subepithelial myofibroblasts known as Foxl1^+^ telocytes. Importantly, either elimination of Foxl1^+^ telocytes or ablation of all Wnt signals emanating from these cells causes rapid crypt failure and death in mice, demonstrating that these cells are a critical component of the intestinal stem cell niche (Aoki et al. 2016; Shoshkes-Carmel et al. 2018; Kondo and Kaestner 2019).

Mice null for Foxl1 exhibit delayed villus formation and a defect in the inhibition of epithelial proliferation in nascent villi, suggesting Foxl1 as an important transcription factor controlling mesenchymal to epithelial signaling (Kaestner et al. 1997). A link to epithelial hedgehog signaling was subsequently established by the identification of functionally relevant binding sites for the hedgehog-dependent Gli transcription factors within an ultra-conserved enhancer at the *Foxl1* locus (Madison et al. 2009). Foxl1 expression was induced in fetal gut mesenchyme explants treated with Shh, while its mRNA levels were reduced in mice deficient for the hedgehog dependent transcription factors Gli2 and Gli3. Taken together, these findings suggested that Foxl1 is a critical mediator of epithelial to mesenchymal cross-talk during villus formation. Here, we set out to determine the molecular targets and pathways controlled by Foxl1 during intestinal villification.

## Results

More than two decades ago, we had reported a developmental delay in *Foxl1* null mice but had focused our analysis on the role of this transcription factor in adult intestinal homeostasis (Kaestner et al. 1997). Using the advanced molecular and imaging technologies available today, we have now re-analyzed the developmental phenotype of these mice. Histological analysis shows that as villus formation is well on its way in wild type fetuses at 15.5 dpc, this process has barely been initiated in *Foxl1* deficient mice (Figure 1A,B). Even two days later, at 17.5 dpc, villification is abnormal in mutant mice, with apparent bridging of nascent villi across the gut lumen, likely reflecting persistent epithelial ridges (Figure 1C, D), a phenotype that persists at 18.5 dpc (Figure 1E,F). In order to investigate this phenotype in three dimensions, we performed scanning electron microscopy. As shown in Figure 1G-L, in control mice villification occurs through the formation of regularly spaced short invaginations into the gut lumen, which by 18.5 dpc have progressed to form elongated villi. This process is dramatically altered in the absence of Foxl1, with absence of the regularly spaced nascent villi and presence instead of long epithelial ridges at 15.5 dpc (yellow arrow in Figure 1H), which persist until late gestation (Figure 1J, red arrow). The presence of epithelial ridges instead of villi is reminiscent of the phenotype seen in small intestinal explants treated with the pan-BMP inhibitor dorsomorphin (Walton *et al*., 2016b), a connection we explore further below.

**Figure 1:**
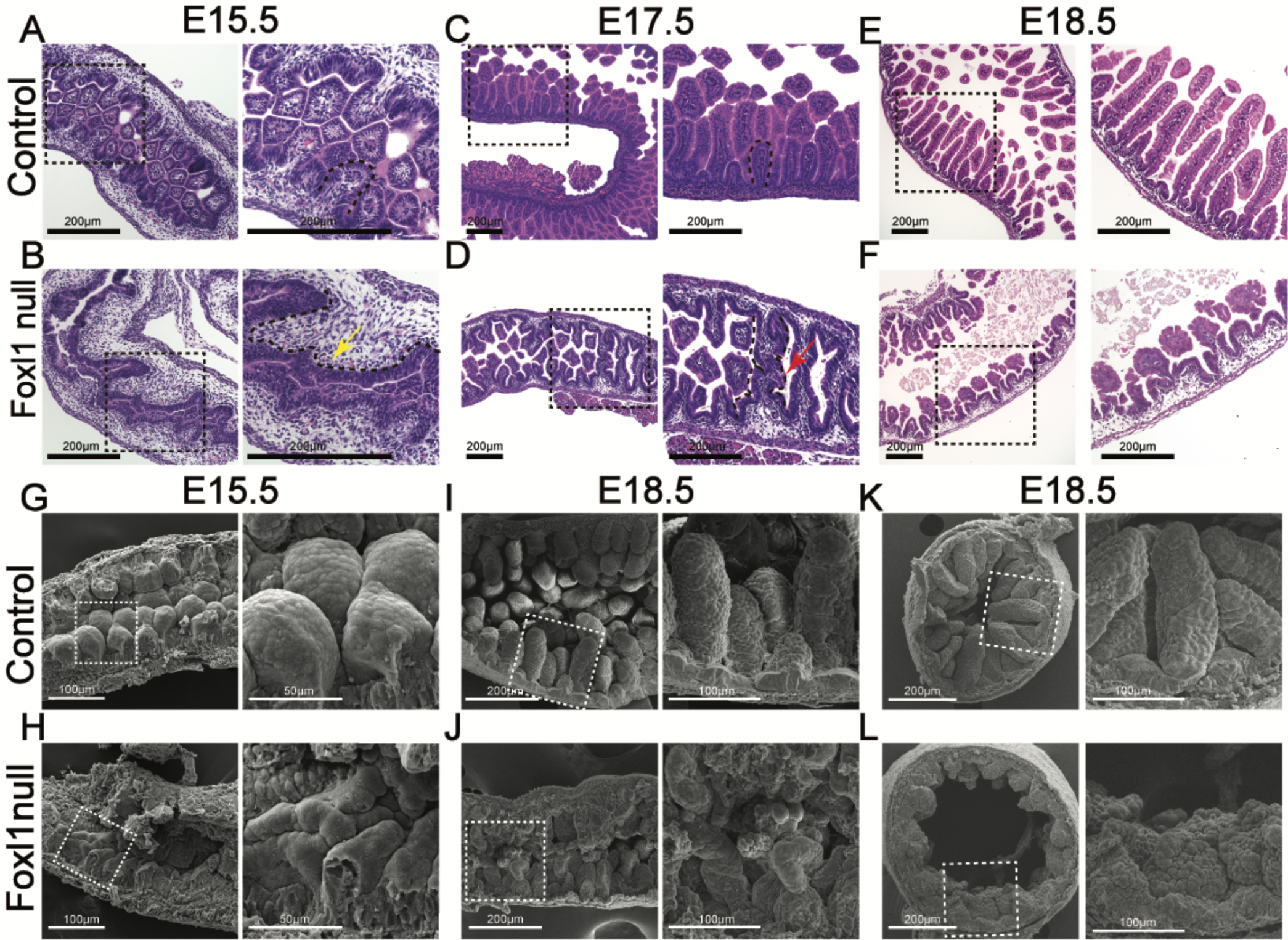
Intestinal villification is dependent on the winged helix transcription factor FOXL1. (A-F) Hematoxylin and eosin stained small intestine from control (A,C,E), or Foxl1 null (B,D,F) fetuses at the developmental stages indicated. The yellow arrow in B indicates lack of invagination, and the red arrow in D marks persistent epithelial ridges. (G-L) Scanning electron micrographs of small intestine from control (G,I,K), or Foxl1 null (H,J,L) fetuses at the developmental stages indicated. In all panels, larger magnification images shown on the right correspond to the areas outlined on the left. Magnification is indicated by scale bars.

### Fetal gut telocytes progenitors exist in two subpopulations

The inclusion of tdTomato in the recently developed *Foxl1* mutant allele employed here (Kolev et al. 2021) enabled us to follow the fate and determine the molecular properties of Foxl1-positive cells embryos heterozygous and homozygous for this Foxl1 null allele. Of note, the *Foxl1* phenotype is recessive, as *Foxl1* heterozygous mice are indistinguishable from wild type controls (Kaestner et al. 1997; Perreault et al. 2001; Katz et al. 2004; Perreault et al. 2005). We performed scRNAseq on the proximal half of the small intestine of 15.5 dpc *FoxL1*^CreER-tdTom/+^ fetuses (Figure 2A,B). As shown in the UMAP analysis in Figure 2C and D, Foxl1^+^ cells segregate into two closely related cell clusters, which we termed ‘telocyte progenitors 1 and 2’. A list of marker genes for all cell populations shown in Figure 2C is given in Supplementary Table 1. Telocyte progenitors 1 express high levels of *Pdgfra* and multiple *Bmp* mRNAs (Figure 1E), and thus most likely correspond to ‘villus cluster cells’, i.e the PDGFRα positive cells important in villus formation originally identified by Karlsson and colleagues (Karlsson et al. 2000). By exclusion, the telocyte progenitor 2 population likely represents Foxl1^+^ cells directly adjacent to the intervillus epithelium, i.e. the presumptive future crypt cells. Interestingly, we identified *Glp2r*, encoding the GLP2 receptor, as another gene predominantly expressed in telocyte progenitor 1 cells, and confirm the localization of the Glp2r mRNA to villus cluster cells by RNAscope analysis (Figure 2F,G). Of note, this single cell analysis clearly shows that Foxl1^+^ cells are distinct from myofibroblast, interstitial cells of Cajal, and pericytes, identified by the expression of Acta2^med^/Myh11^med^/Des^med^/Tagln^med^, Etv1^+^/Kit^+^/Acta^low^, and Cspg4^+^/Pdgfrb^+^/Abcc9^+^ or Abcc9^+^/Ndufa4l2^+^, respectively. The two Foxl1^+^ telocytes populations are most closely related to a Foxl1-negative mesodermal populations. Future studies will be directed towards exploring if a progenitor/descendant relationship exists between these cell types. Major hallmarks of the telocyte progenitor 1 and 2 populations are summarized in the graphic shown in panel 2H.

**Figure 2:**
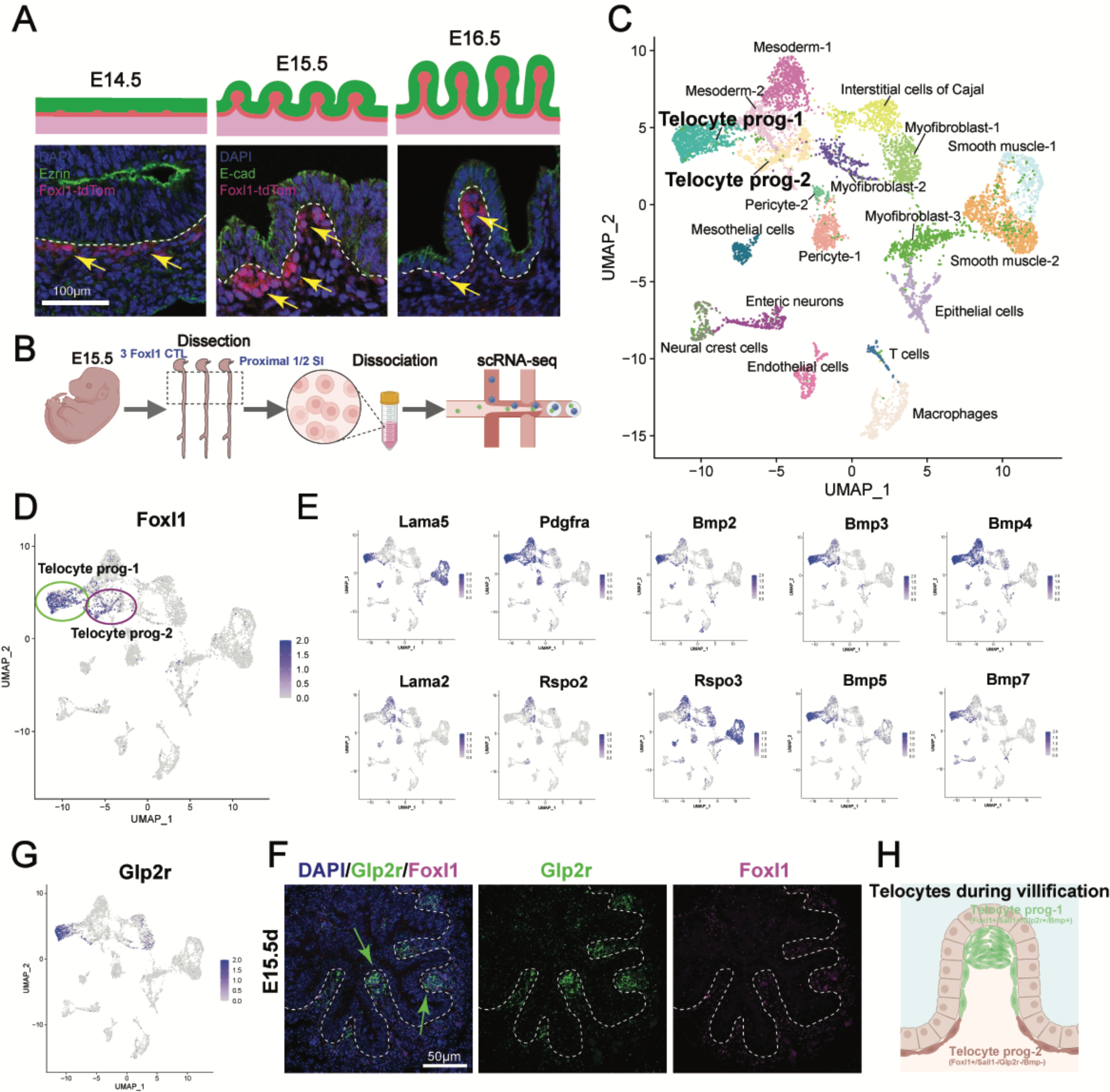
Fetal FOXL1-positive telocyte progenitors partition into villus cluster and intervillus populations. (A) FOXL1-positive cells labeled by tdTomato expression restructure during the epithelial transition, when the pseudostratified epithelium (E14.5) is converted to a simple cuboidal epithelium (E16.5). Epithelial cells are labeled with Ezrin or E-cadherin (E-cad) in green, Foxl1-positive cells by tdTomato in red). (B) Experimental outline for single cell RNAseq study. The proximal half of the E15.5 small intestine was used for the analysis. (C) UMAP plot of scRNAseq data identifies more than a dozen cell types. Foxl1-positive cells are labeled as telocyte progenitors 1 and 2. (D) UMAP plot showing mRNA levels of Foxl1. (E) UMAP plot of mRNA expression of multiple genes that differentiate between the two telocyte progenitor populations. Note the high expression of multiple BMP mRNAs in telocyte progenitors (F) UMAP plot of Glp2r, encoding the receptor for the intestinotrophic hormone GLP-2. Glp2r expression is highly enriched in telocyte progenitor 1 cells. (G) RNAscope analysis to detect Glp2r transcripts (green) and Foxl1 mRNA (purple). The Glp2r mRNA is highly enriched in villus tip telocyte progenitors. (H) Model of relative positioning of telocyte progenitors 1 and 2 during intestinal villification.

### Expression of PDGFRα in telocyte progenitors is dependent on Foxl1

Next, we analyzed the expression profiles of the fetal small intestine of Foxl1 null (*FoxL1*^CreER-tdTom/CreER-tdTom^) mice and compared them to those of heterozygous fetuses (Figure 3A). Note that in both control and Foxl1 null fetuses, Foxl1-expressing cells are localized to the mesodermal cell layer that is directly juxtaposed to the developing epithelium, and loss of Foxl1 does not affect either telocyte progenitor number or position (Figure 3B). The UMAP plot shown in Figure 3C clearly demonstrates that the telocyte progenitor 2 population is dramatically reduced in abundance in the Foxl1 null intestine. As mentioned above, among the markers of the telocyte progenitor 1 cells is PDGFRα. The scRNAseq data shown in Figure 3D as well as the immunofluorescence staining presented in Figure 3E demonstrate the PDGFRα expression is Foxl1 dependent. To determine if *Pdgfra* is a direct target of Foxl1, we performed Cut-and-Run assays on telocytes from 15.5 dpc fetal gut. However, we found no Foxl1 binding event within 10 kb of the *Pdgfra* promoter.

**Figure 3:**
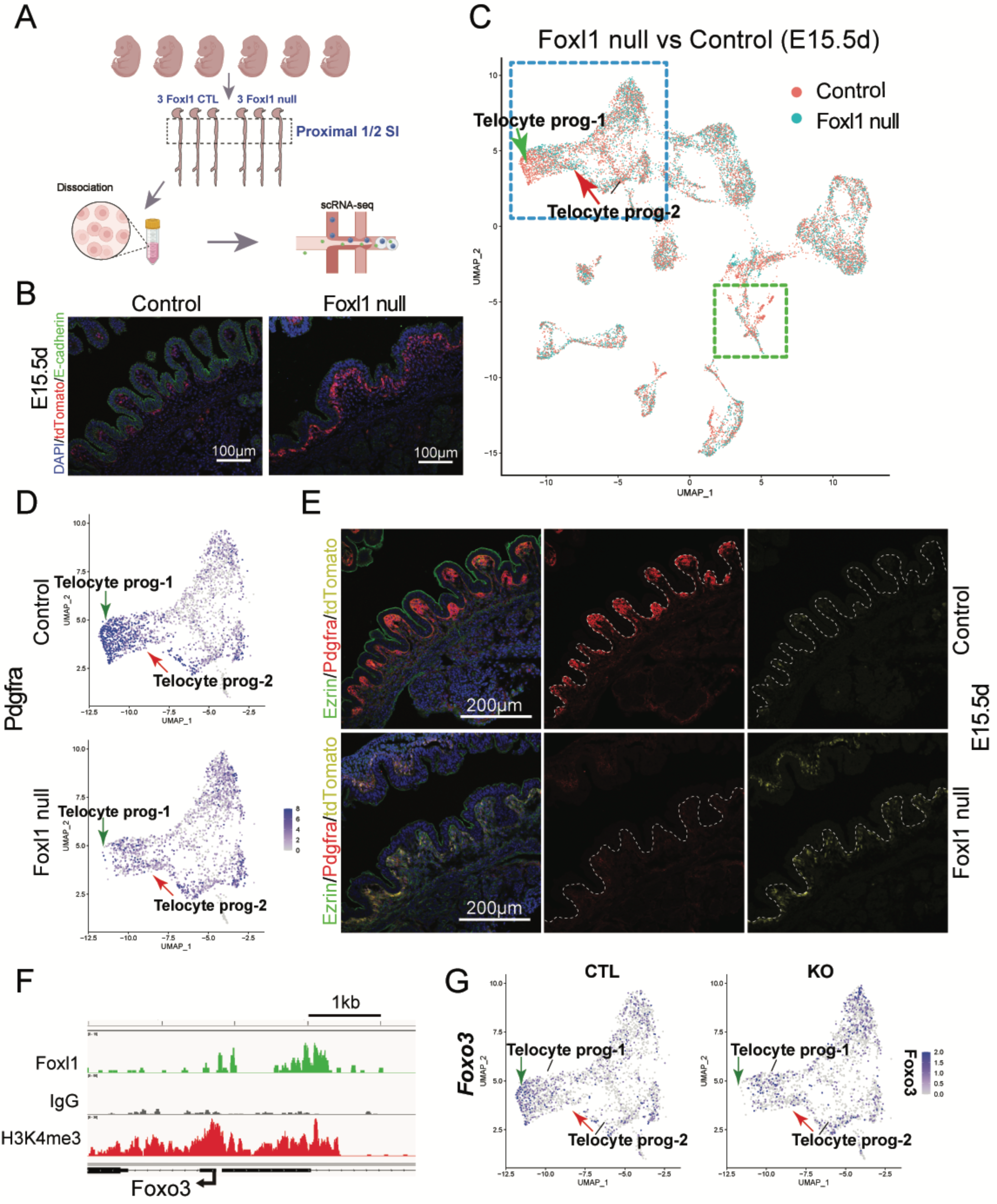
Villus cluster PDGFRα expression is Foxl1 dependent. (A) Experimental outline for the determination of Foxl1-dependent gene expression by scRNAseq. (B) Foxl1-positive telocytes, labeled by tdTomato expression (red) are retained in number and position in Foxl1 null mice. Epithelial cells are labeled by E-cadherin (green). (C) UMAP plot of scRNAseq data shows dramatic shifts in both telocyte progenitor (blue box) as well as in epithelial cells (green box). Single cells are labeled by genotype. (D) UMAP plot showing PDGFRa expression in control and Foxl1 null telocyte progenitors. (E) Immunofluorescence labeling confirms dramatic reduction in PDGFRα expression (red) in the absence of Foxl1. The apical membrane of epithelial cells is stained for Ezrin (green). The persistence of telocyte progenitors in *Foxl1* null mice is confirmed by staining for the tdTomato transgene (yellow) which replaces the *Foxl1* coding region in this *Foxl1* null allele. (F) Cut and Run analysis of fetal telocytes indicates that the *Foxo3* promoter is in an active state, as indicated by a strong H3K4me3 peak (red trace). Foxl1 is bound to the proximal promoter of *Foxo3*. (G) Foxo3 expression in fetal telocyte progenitors is Foxl1-dependent as shown by reduced expression levels in the Foxl1 null intestine by scRNAseq analysis.

It was previously reported that the related winged helix transcription factor Foxo3 is a transcriptional activator of *Pdgfra,* which binds to an evolutionarily conserved *cis*-regulatory element in its proximal promoter (Mei et al., 2012). Therefore, we hypothesized that Foxl1 might indirectly control *Pdgfra* via activation of *Foxo3*. Using Cut-and-Run of fetal telocytes, we indeed found several Foxl1 binding sites in the *Foxo3* promoter (Figure 3F). In addition, our scRNAseq data show a reduction of Foxo3 transcript levels in the telocyte progenitor 1 population of Foxl1 null fetuses (Figure 3G).

### Foxl1 controls multiple Bmp genes in telocyte progenitor cells

Next, we focused on BMP proteins, as BMP signaling from the mesoderm to the epithelium is critical for villus cluster formation (Walton et al. 2016b). Telocyte-produced BMPs enriched in the villus cluster cells signal to the overlaying endoderm to inhibit Wnt signaling and limit proliferation (Figure 4A). High expression of several BMP genes was present in particular in telocyte-progenitor 1 cells (Figure 4B). In case of BMP4, expression also extends to Foxl1 negative GPX3+ FLCs; however, its levels are clearly reduced specifically in Foxl1-deficient telocytes (Figure 4C). Using our Cut-and-Run data from sorted 15.5 dpc telocytes, we identified a Foxl1 binding site ∼55kb downstream of the *Bmp4* promoter, suggesting direct activation by Foxl1 (Figure 4C). Loss of mesenchymal BMP signals is expected to result in de-inhibition of Wnt signaling in the epithelium overlying the villus cluster telocytes. Indeed, we found expression of the Wnt target Sox9 expanded from the developing crypts to nascent villi in the Foxl1 null fetal intestine (Figure 4D). Likewise, epithelial proliferation was not confined to the nascent crypts but extended to the villus epithelium in the mutant mice (Figure 4D). Thus, Foxl1 is a critical factor required for the demarcation of the postmitotic villus from the mitotic intervillus epithelium.

**Figure 4:**
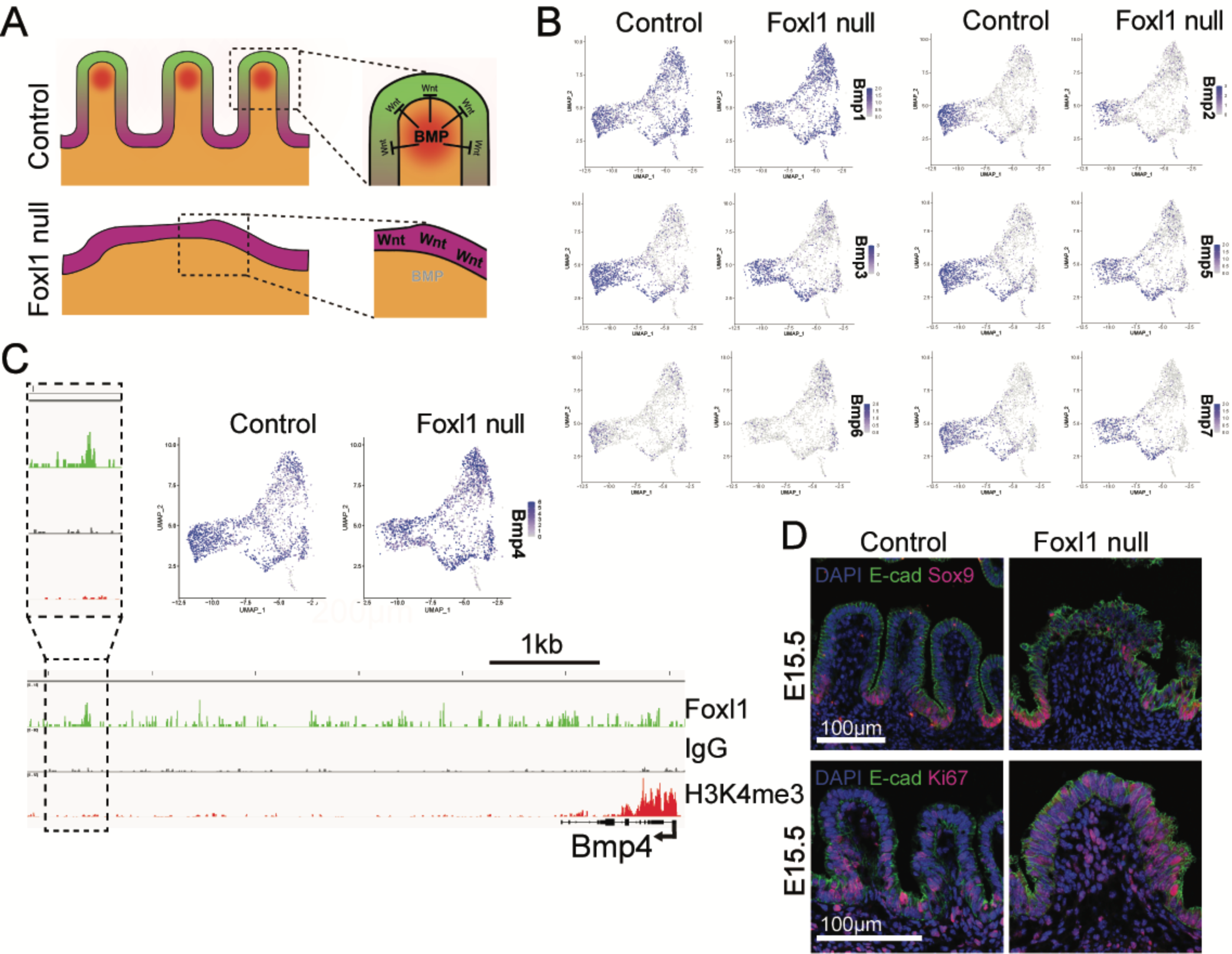
Loss of BMP expression and abnormal epithelial proliferation in the Foxl1-deficient fetal small intestine. (A) Top: model for partitioning of small intestinal epithelial cells into postmitotic villus cells and proliferating intervillus cells through BMP signaling emanating from villus cluster cells. Bottom: In absence of Foxl1, BMP signals are reduced, and Wnt dependent proliferation is maintained in a larger epithelial domain. (B) Expression of multiple Bmp genes is reduced in telocyte progenitor 1 cells in absence of Foxl1. (C) Bmp4 expression is reduced in absence of Foxl1, and Foxl1 binds downstream of the Bmp4 gene by Cut and Run assay of fetal telocyte progenitors. (D) Expression of the Wnt target gene Sox9 (purple) is restricted to intervillus regions in the E15.5 control small intestine, but maintained in a subset of villus epithelial cells in Foxl1 null mice. Likewise, proliferating epithelial cells, marked by Ki67 (purple) become restricted to intervillus regions in control mice, but extend over the emerging villus tip in absence of Foxl1. Epithelial cells are marked by E-cadherin (green).

### Foxl1 is required for mesenchymal expression of planar cell polarity genes

Recently, planar cell polarity genes were identified among the mesenchymal Gli targets in the fetal gut, and it was demonstrated further that the GLI2 target *Fat4* is required for villus development during the epithelial transition (Rao-Bhatia et al. 2020). The discovery that the PCP pathway acts within the mesenchymal compartment to structure stromal cells was surprising, as typically PCP pathway function has been reported within epithelial cell layers (Butler and Wallingford 2017). As documented above, before the epithelial transition, Foxl1+ cells form a uniform cell layer with a depth of only one to two cells surrounding the primitive gut tube, which is then patterned into the telocyte progenitor 1 (villus cluster) and telocyte progenitor 2 (crypt base) cells, possibly with the involvement of the PCP system. We found several PCP genes (*Fat4*, *Wnt5a*, *Vangl1* and *2*) that exhibit reduced expression in the absence of Foxl1 (Figure 5A). To evaluate if planar cell polarity is affected by the absence of Foxl1, we stained small intestinal sections from 15.5 dpc fetuses with E-cadherin, which marks the adherens junctions of epithelial cells. While in the wild-type intestine, epithelial cells had already arranged into a simple cuboidal epithelium with regularly spaced junctional complexes, the epithelium was disorganized and partially multilayered in the absence of Foxl1 (Figure 5B).

**Figure 5:**
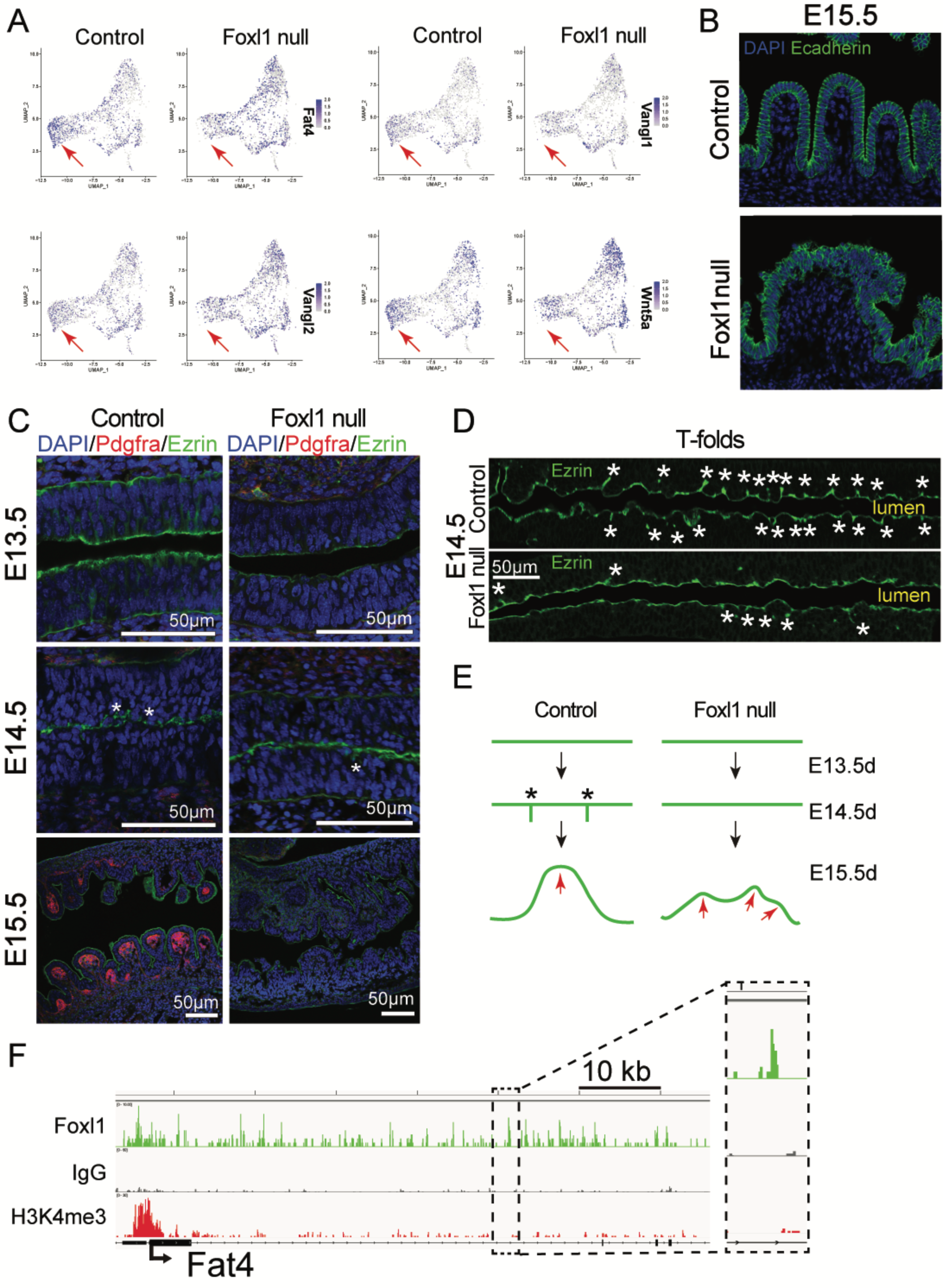
Foxl1 is required for full activation of planar cell polarity genes in small intestinal telocyte progenitors. (A) UMAP plots show reduced expression of planar cell polarity genes Fat4, Vangl1, Vangl2, and Wnt5a in Foxl1-deficient telocyte progenitors. (B) E-cadherin staining (green) shows ordered cuboidal epithelial cells in the control fetal intestine, while epithelial structure is perturbed and partially multilayered in the absence of Foxl1. (C) Staining of control fetal small intestine at time-points flanking the epithelial transition with Ezrin (green) and PDGFRα (red) shows the emergence of T-folds at E14.5 and PDGFRα positive villus clusters by E15.5. In *Foxl1* null mice, formation of T-folds is delayed. (D) Detail of T-fold formation as revealed by Ezrin staining. (E) Model for villification in control and Foxl1 null mice. (F) Cut and Run analysis shows binding of Foxl1 at the *Fat4* promoter and downstream elements.

Rao-Bhatia and colleagues had found that villification defects in mice with mutations in the PCP gene Fat4, which were worsened by simultaneous heterozygous loss of Vangl2 (Rao-Bhatia et al. 2020). This defect was accompanied by a reduction in the number of epithelial T-folds, characteristic invaginations of the epithelium that can be visualized by staining with the apical membrane marker Ezrin that form the boundaries of developing villi. In order to evaluate if the reduced expression of PCP genes in *Foxl1* null mice impacts epithelial remodeling, we stained small intestinal sections from fetuses at developmental stages spanning villification (13.5 to 15.5 dpc) for Ezrin to identify T-folds and PDGFRα to label villus cluster cells. As shown in Figure 5 C-E, the number of T-folds is clearly reduced in the Foxl1 deficient intestine, coinciding with the loss of PDGFRα expression in telocyte progenitors. Of note, Cut-and-Run analysis of transcription factor occupancy showed Foxl1 binding both at the promoter and at a distal downstream enhancer of *Fat4*, indicating a direct regulatory relationship (Figure 5F).

### Loss of Foxl1 impacts epithelial gene expression profiles

As shown above, Foxl1 deficiency impacts the patterning of the overlying epithelium, with most villus tip epithelial cells remaining in the cell cycle (Figure 4D). We had also noted a shift in the UMAP pattern of epithelial cells between control and Foxl1 null cells in 15.5 dpc embryos (Green box in Figure 3C). To address this issue further, we reclustered epithelial cells only via UMAP. Figure 6A shows the fetal gut epithelial cells of control embryos partition into two major groups, which we identified as ‘secretory progenitors’ and ‘undifferentiated epithelial cells’ based on their expression profile. The heatmap in Figure 6B shows the 225 most differentially expressed genes between these two clusters (false discovery rate < 10%; absolute fold-change >2), while Figure 6C indicates selected markers genes for each cell type. Fetal secretory progenitors are characterized by high levels of the mRNAs for transcription factors *Klf4*, *Spdef*, and *Sox4*, known to be critical for secretory cell differentiation (Katz et al. 2002; Gregorieff et al. 2009; Gracz et al. 2018). Undifferentiated epithelial cells in contrast exhibit strong expression of Sox9 (which marks them as proliferating intervilllus cells as seen in Figure 4D) as well as markers of the absorptive enterocyte lineage (*Alpi*, the gene for intestinal alkaline phosphatase, *Fabp1*, encoding fatty acid binding protein 1, *Apoa4* the gene for Apolipoprotein A4 which is important in intestinal cholesterol absorption, *Slc16a1*, encoding the monocarboxylic acid transporter for lactate).

**Figure 6:**
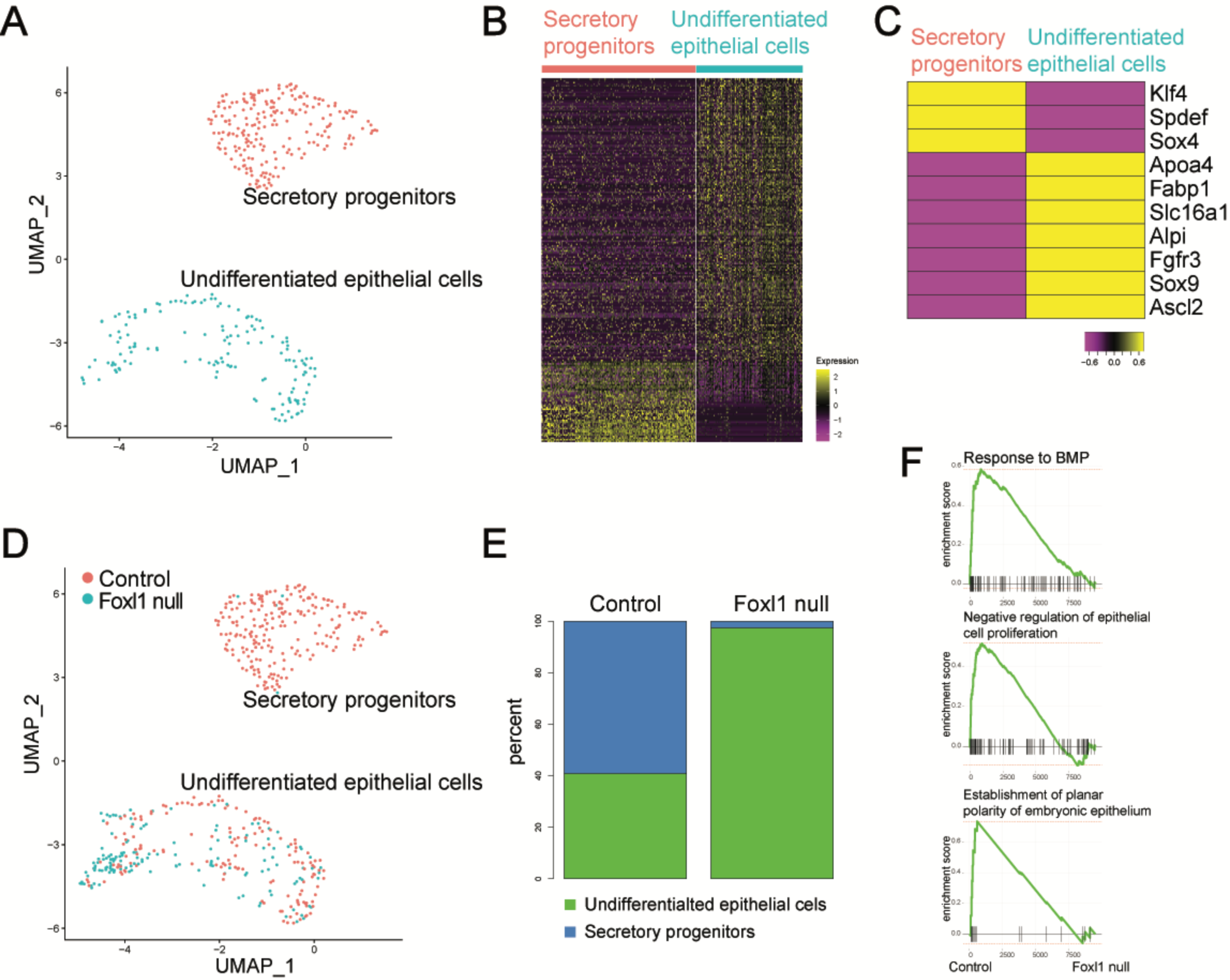
Foxl1-dependent telocyte signals are required for epithelial gene expression. (A) UMAP plot of control epithelial cells from E15.5 embryos. The cells partition into two major clusters of secretory progenitors (red) and undifferentiated epithelial cells (blue). (B) Heatmap of genes differentially expressed between secretory progenitors and undifferentiated epithelial cells. Each row represents a gene, each column a cell. (C) Expression of key marker genes in secretory progenitors and undifferentiated epithelial cells (D) UMAP plot of control (red) and Foxl1 null (blue) epithelial cells. Note that only very few Foxl1 null cells cluster with secretory progenitor cells. (E) Proportion of secretory progenitors and undifferentiated epithelial cells in control and Foxl1 null embryos. (F) Gene set enrichment analysis (GSEA) identifies critical pathways as Foxl1 dependent. The normalize enrichment scores (NES) were 2.18 for the ‘response to BMP’, 1.97 for ‘negative regulation of epithelial proliferation’, and 1.90 for ‘establishment of planar polarity of embryonic epithelium.

Next, we added epithelial cells from Foxl1 null embryos to the UMAP, and found that they are largely confined to the undifferentiated epithelial cell cluster (Figure 6D). When we quantified the proportion of cells in each cluster, we found a striking loss of secretory progenitor cells in the Foxl1 null embryos (Figure 6E). Finally, we performed gene set enrichment analysis to search for pathways that are differentially regulated in the absence of Foxl1. As shown in Figure 6F, the response to BMP signaling, negative regulation of epithelial proliferation, and establishment of planar cell polarity were all strongly enriched among the genes more highly expressed in the control gut epithelium, confirming that loss of telocyte Foxl1 has a major impact on the development of the fetal intestinal epithelium.

## Discussion

Villus formation is a fascinating and obviously essential process in vertebrate gut development and depends on reciprocal epithelial-to-mesenchymal cross talk. Epithelial hedgehog proteins are among the earliest signals emanating from the endoderm during organogenesis of the gut. Consequently, inhibition of hedgehog signaling with neutralizing antibodies or ablation of either *Shh* (sonic hedgehog) or *Ihh* (Indian hedgehog) causes impairment of gut development and villus formation (Motoyama et al. 1998; Ramalho-Santos et al. 2000). Hedgehog proteins signal via their receptor Ptch1 (patched 1), expressed exclusively in the gut mesoderm, to stabilize the DNA-binding transcription factors Gli2 and Gli3. In 2006, computational analysis of evolutionarily conserved enhancers led to the identification of an ultra-conserved putative enhancer located between the neighboring *Foxl1* and *Foxf1* genes (Hallikas et al. 2006). We identified seven Gli binding sites in this genomic region, some conserved from Fugu to human, and showed through *in vitro* and *in vivo* studies that both *Foxl1* and *Foxf1* are Gli target genes (Madison et al. 2009). Here, we demonstrate that Foxl1-expressing telocyte progenitors are partitioned into two major subpopulations with distinct gene expression profiles. Telocyte progenitors 1 and 2 correspond to telocytes in the villus clusters and those adjacent to developing crypts, respectively.

Villification is strongly impaired in the absence of *Foxl1*, and abnormal proliferation persist in epithelial cells in the developing villi. We attribute this to the loss of BMP signaling as multiple *Bmp* genes exhibit reduced expression in absence of Foxl1. BMP signaling had been established by Walton and colleagues as key factor of villus formation (Walton et al. 2016b). Recently, planar cell polarity genes were identified among the mesenchymal GLI2 targets in the fetal gut, and it was demonstrated further that the GLI2 target *Fat4* is required for villus development during the epithelial transition (Rao-Bhatia et al. 2020). The discovery that the PCP pathway acts within the mesenchymal compartment to structure stromal cells is novel and exciting, as typically PCP pathway function had been reported within epithelial cell layers (Butler and Wallingford 2017). The model of villification proposed by Rao-Bhatia and colleagues states that activation of GLI2 in telocyte progenitors opposite to the hedgehog-secreting epithelium is sufficient to activate *Fat4* and other planar cell polarity genes directly (Rao-Bhatia et al. 2020). Our data indicate that PCP induction in the developing gut mesenchyme is more complex than proposed by Rao-Bhatia and colleagues and depends not only on GLI proteins but also on Fox transcription factors active in telocyte progenitors. We document by scRNAseq analysis of the proximal small intestine of 15.5 dpc fetuses that expression of planar cell polarity genes including *Fat4* is enriched in Foxl1+ telocyte progenitors and reduced dramatically in the absence of Foxl1 (Figure 5). Therefore, we propose a feed-forward loop for the regulation of stromal planar cell polarity genes, in which GLI2 activates both *Foxl1* and *Fat4* (and related targets), while Foxl1 also activates planar cell polarity gene expression in epithelium-adjacent stromal cells in a coherent feed-forward loop (Figure 7). Loss of the telocyte transcription factor Foxl1 has a major secondary impact on the overlying epithelium, were reduced BMP signaling lead to loss of secretory progenitor differentiation and retention of proliferating epithelial cells overlying villus cluster cells.

**Figure 7:**
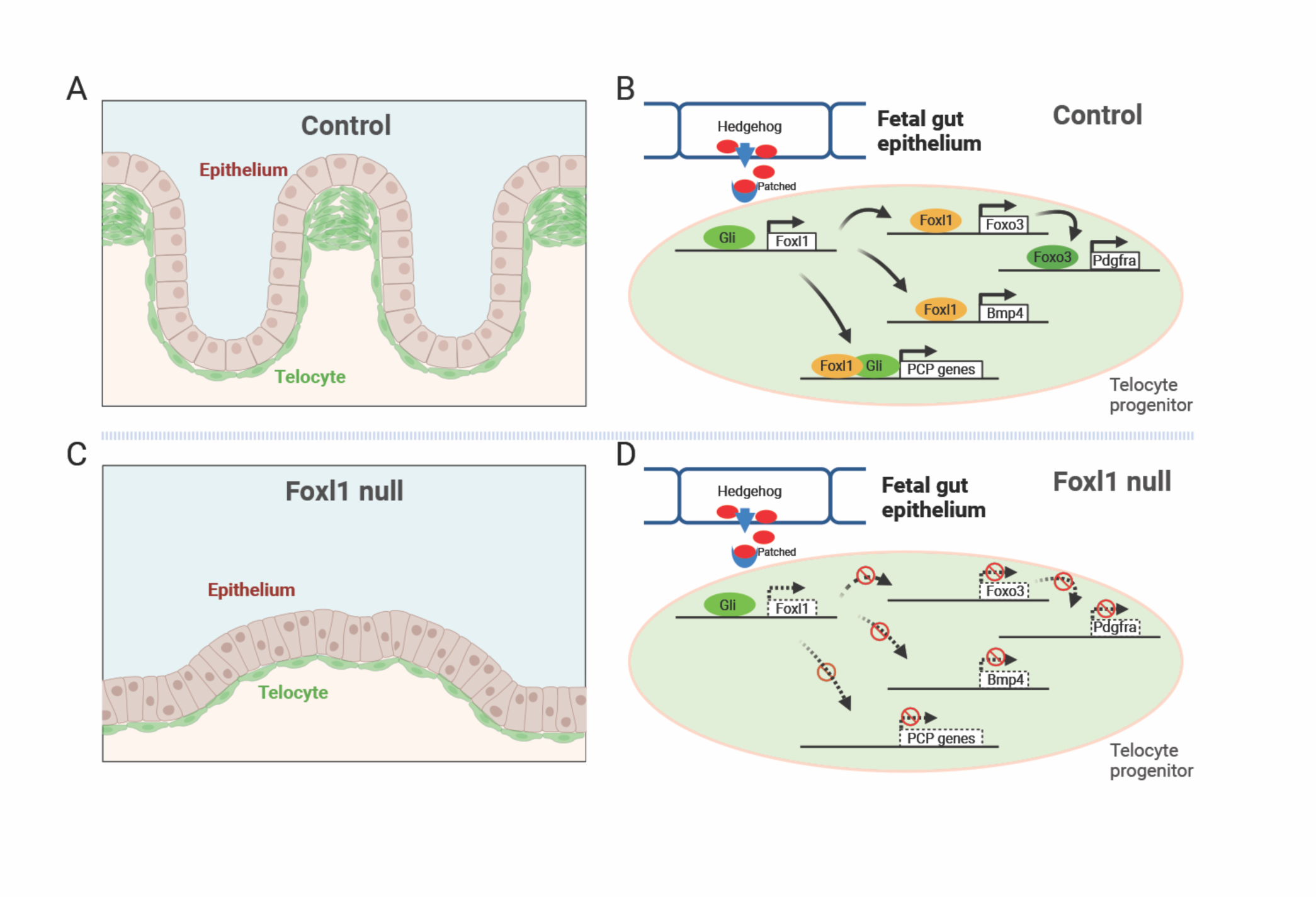
Stromal Foxl1 is required for villification of the fetal intestine. (A) Foxl1-positive telocytes are partitioned into villus cluster and intervillus cells in the control small intestine. (B) Hedgehog signals from the overla

Another interesting telocyte gene we identified by our single cell RNAseq analysis is *Glp2r*, encoding the receptor for the intestinotrophic hormone GLP-2. Expression of Glp2r is maintained in adult telocytes (data not shown). GLP-2 is secreted by intestinal L cells, likely in response to luminal glucose in the ileum and colon. While in the healthy gut all glucose is absorbed in the duodenum and jejunum, when small intestinal absorptive capacity is limited such is the case after small intestinal injury, residual glucose enters the distal gut, triggering GLP-2 release, which then signals to the upper small intestine to increase crypt proliferation via production of growth factors such as IGF1 and EGF in the GLP-2 responsive cells (Dube et al. 2006; Fesler et al. 2020). The identity of the GLP-2-responsive cell has been controversial because of the lack of specific antibodies (Yusta et al. 2019). Telocytes are the logical site for *Glp2r* expression, as they are only separated from the crypt epithelium by the 60 nm basement membrane (Shoshkes-Carmel et al. 2018). Future studies employing telocyte-specific ablation of *Glp2r* will address whether GLP2 signaling is indeed mediated by Foxl1-positive cells.

In conclusion, we have shown that the winged helix transcription factor, expressed in endoderm-adjacent mesodermal telocytes, is critical for the epithelial transition, epithelial gene expression, and ordered villus formation via the regulation of BMP, PDGFRa, and PCP signaling molecules.

## Materials and Methods

### Mice

All animal procedures were approved by Institutional Animal Care and Use Committee of the Office of Animal Welfare at the University of Pennsylvania and conducted under protocol 804436. The mice used in this study were housed in a Specific Pathogen Free (SPF) facility at the University of Pennsylvania’s animal center. They were individually housed in ventilated cages and provided with controlled temperature, humidity, a 12-hour light-dark cycle, a standard rodent chow diet, and constant access to water.C57BL/6 wild-type mice were obtained from Jackson Laboratory (Stock number: 000664). The Foxl1CreERT2-TdTomato gene replacement allele was described previously (Kolev et al. 2021).

For embryonic studies, embryonic day 0.5 (0.5 dpc) was defined as noon on the day when the copulatory plug was observed. Since homozygotes of the Foxl1CreERT2-TdTomato gene replacement allele are viable and fertile in adulthood, the Foxl1 null embryos were generated by either crossing two heterozygotes or crossing one homozygote with a heterozygote. Single cells for the Cut & Run experiment were isolated from dissected embryonic small intestines at 15.5 dpc, which were obtained by crossing Foxl1CreERT2-tdTomato heterozygous mice with C57BL/6 wild-type mice.

Embryonic data were collected from developmental stages before (13.5 dpc), during (14.5 dpc), and after (15.5-18.5 dpc) the onset of villification in the developing small intestines. Due to the indistinguishable sex of the embryos and the absence of reported sex differences in villus morphogenesis, mice of both sexes were included in all experimental procedures. Therefore, random assignment of mice from either sex was conducted within each experimental group.

### Tissue isolation and sectioning

Mouse embryonic small intestinal samples from different stages were carefully dissected and harvested from the body. All samples used for histological sectioning and subsequent staining were taken from duodenal sections. For paraffin section, samples were fixed overnight at 4° C temperature in 4% paraformaldehyde in PBS buffer (Invitrogen) and incubated in 70% ethanol solution after three times’ wash with PBS buffer for paraffin section, and then were submitted to the Molecular Pathology and Imaging Core of the Center for Molecular Studies in Digestive and Liver Diseases (P30 DK050306) for further process, embed and sectioning for sectioning at tum thickness and then sectioned slices were kept at room temperature. For frozen section, samples were incubated overnight in 30% sucrose in PBS buffer after overnighted fixation at 4°C in 4% paraformaldehyde in PBS buffer (Invitrogen) until the intestinal tissues completely sink to the tube bottom, and then were embed in OCT for quick-frozen and stored at -80°C. OCT-embedded samples were performed the cryo-sectioning though a using a cryostat (Cryostar NX50, ThermoFisher Scientific) at 10 µm, dried for 10 minutes at room temperature before the further immunofluorescent staining or stored at - 80°C.

### Scanning Electron Microscopy

The scanning electron microscope (SEM) experiments were conducted at the microscopy core facility of the UPenn Department of Cell and Developmental Biology. After dissection, the presumptive duodenal sections of the 15.5 and 18.5 dpc fetuses were washed three times with 50 mM Na-cacodylate buffer and fixed overnight using a solution consisting of 2.5% glutaraldehyde in 50 mM Na-cacodylate buffer at a pH of 7.3. Subsequently, samples were dehydrated using a graded series of ethanol, gradually reaching 100% ethanol over a span of 1.5 hours. After dehydration, samples were incubated for 20 minutes in a solution containing 50% HMDS (Sigma-Aldrich) in ethanol, followed by three changes of 100% HMDS. After air-drying overnight, samples were mounted on stubs and coated with a layer of gold palladium using the sputter coating technique. Finally, we observed and photographed the specimens utilizing a Quanta 250 FEG scanning electron microscope manufactured by FEI (Hillsboro, OR, USA) with a 10 kV accelerating voltage.

### Histology and Immunofluorescence

Paraffin sections were deparaffinized and rehydrated using xylene and descending ethanol gradients. For H&E staining, tissues were stained with Harris’ Hematoxylin and alcoholic Eosin Y. For IF staining, paraffin sections were subjected to antigen retrieval. For the frozen slices, OCT was directly removed in sterile water. Then the slices were performed the incubation with primary antibodies overnight at 4 °C after serum blocking for 1 hour at room temperature and then with appropriate secondary antibodies at room temperature in the dark for 1 hour.

The following antibodies were used: goat anti-PDGFR alpha 1: 200 (R&D Systems, AF1062), mouseanti-EZRIN 1:1000 (Sigma-Aldrich, E8897), rat anti-E-Cadherin antibody [DECMA-1] 1: 200 (Abcam, ab11512), goat anti-E-cadherin 1: 200 (R&D systems, AF-648), rabbit anti-Sox9 1: 200 (Abcam, ab185966), rabbit anti-Ki67 1:200 (Abcam, ab16667), and Alexa Fluor 488-, 594- and 647-conjugated secondary antibodies, obtained from Life Technologies. Then, slides were stained with DAPI (Sigma-Aldrich, D9542), sealed and photographed using a confocal microscope Leica Stellaris 5. The generated confocal images were further edited using Photoshop and Adobe Illustrator.

### Single-cell capturing and cDNA library construction

Timed embryos were obtained by crossing homozygous male mice with heterozygous female mice. The small intestine was obtained through dissection, and half of the portion close to the stomach was collected after folding it in half. Additionally, the tail of each embryo was collected for genomic DNA extraction using the KAPA Mouse Genotyping Kit HotStart (Kapa Biosystems, KK7352). Subsequently, genotyping was performed to distinguish between heterozygotes and homozygotes.

Isolated intestinal cells were digested with collagenase type II and DNase I to prepare the single-cell suspension and then sorted in phosphate-buffered saline with 0.05% BSA to enrich the living cells through FACS sorting (MoFlo Astrios Sorter). Then the cells obtained were measured for cell concentration and viability with Trypan blue using a Countess II Automated Cell Counter from Life Technologies. Then the single-cell suspension was diluted to appropriate concentration and loaded on a 10x Genomics Chromium Single Cell Controller (Pleasanton, CA) with a target of about 5,000 cells per sample. Single-cell library preparation was completed using the 10x Genomics Chromium Single Cell 3’ Library & Gel Bead Kit v2 strictly following manufacturer’s protocol. The quality and quantity testing of obtained short cDNA fragment libraries using an Agilent 2100 Bioanalyzer and Invitrogen Qubit Fluorometer. Finally, the single-cell cDNA Libraries were sequenced on an Illumina Novaseq 6000 instrument.

### Cut & Run and DNA product sequencing

Timed embryos were obtained by crossing homozygous or heterozygous male mice with C57BL/6 female mice. The entire length of the small intestine was digested with collagenase type II and DNase I to reach the single-cell state. Subsequently, the cells were sorted in phosphate-buffered saline with 0.05% BSA to enrich the tdTomato-positive cells using FACS (MoFlo Astrios Sorter).

Isolated tdTomato-positive telocytes from the embryonic small intestines at 15.5 dpc were subjected to CUT&RUN experiments. CUT&RUN experiments were performed using the CUT&RUN Assay Kit (EpiCypher, Catalog No. 14-1048) following the manufacturer’s Instructions using 40,000 telocyte progenitors. Anti-Foxl1, H3K4me3 positive control and rabbit IgG negative control antibodies (13-0042k) were used in these experiments. Purified CUT&RUN DNA products were subjected to the CUT&RUN Library Prep Kit (EpiCypher, Catalog No. 14-1002) for library construction, and the libraries sequenced on an Illumina Hiseq X Ten instrument.

### Single-molecule RNA fluorescent in situ hybridization

Proximal small intestinal sections from the Foxl1 null mice and controls from the same litter were fixed and gradient dehydrated following the methods described above. To perform the single-molecule RNA fluorescence in situ hybridization (smFISH) we employed the RNAScope Multiplex Fluorescent Reagent Kit v2 (323270) from Advanced Cell Diagnostics following the manufacturer’s recommendations. smFISH imaging was performed using a confocal Leica Stellaris 5 microscope. The following RNAscope probes were obtained from Advanced Cell Diagnostics: Foxl1(C3), and Glp2r (C1).

## Acknowledgements

We thank members of the Kaestner lab for helpful discussions and Mark Tigue for maintaining our mouse colony. This work was supported by NIH grants R37DK053839 and R01DK139049. We thank the UPenn Center for Molecular Studies in Digestive and Liver Diseases (P30 DK050306) for the use of the Molecular Pathology and Imaging Core (MPIC) for tissue processing, the UPenn Diabetes Research Center Functional Genomics Core (P30 DK019125) for help with data analysis, and the Cell & Developmental Biology Microscopy Core for the use of their confocal imaging services.

## Author Contribut1ions

GZ and KHK – Conceptualization and writing. GZ, HDM, and DL – Methodology. GZ and JS - Data curation and visualization. KHK - Supervision. KHK – Funding acquisition.

## Supplemental information

Legends for supplementary figures and tables

**Figure S1:**
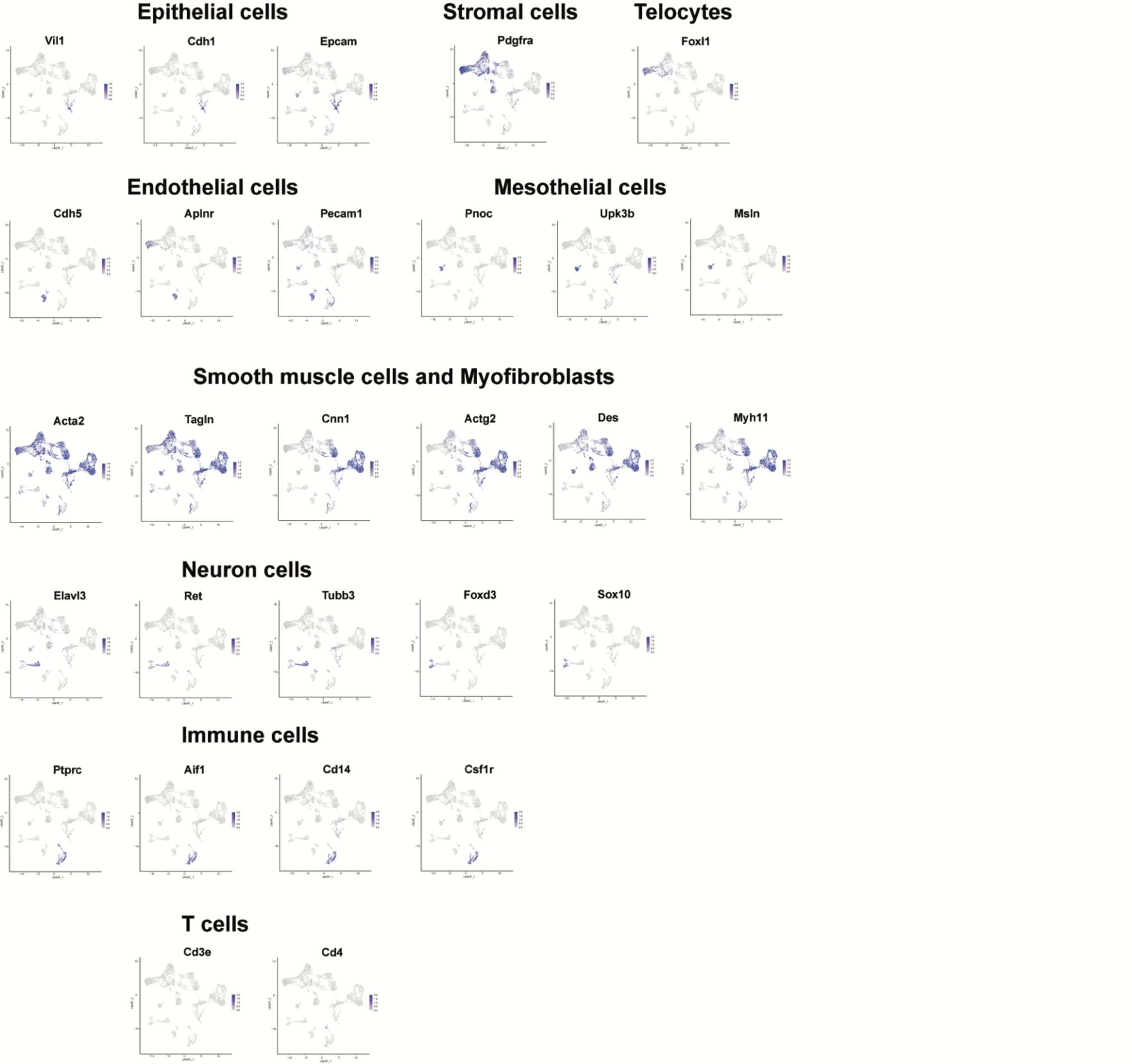
Single nuclei RNA-sequencing of adult mouse colon. (A) A brief description of sample processing and data analysis. Colon tissue from two control mice were used to generate marker genes of the cell types indicated.

